# Assessing the social impact of an online course on the clinical management of Cutaneous Leishmaniasis: A qualitative study

**DOI:** 10.1101/631978

**Authors:** Carme Carrion, Marta Aymerich, Liliana Arroyo, José A Ruiz-Postigo

**Affiliations:** eHealth Center (eHC) and Health Sciences Faculty, Universitat Oberta de Catalunya (UOC), Av. Tibidabo, 39-43, 08035 Barcelona, Catalonia, Spain; Institute for Social Innovation, Dept of Social Sciences, ESADE Business & Law School. Avda Pedralbes, 60-62, 08034 Barcelona, Catalonia, Spain; Leishmaniasis Control Programme, Innovative and Intensified Disease Management Unit, Department of Neglected Tropical Diseases, World Health Organization (WHO), Geneva, Switzerland

## Abstract

**Background:** In conjunction with the World Health Organization (WHO), Universitat Oberta de Catalunya (UOC) has implemented training in the treatment of cutaneous leishmaniasis (CL) for health professionals in countries where the condition is highly endemic through an interactive on-line course. With three editions of the course successfully completed, we consider assessment of the course’s social impact through analysis of the outcomes beyond knowledge acquisition to be of paramount importance.

**Methodology/Principal findings:** To this end, we have conducted a study using the Theory of Change theoretical approach, combined with an exploration of students’ experiences through in-depth interviews before, during and after taking the course. These testimonies have been analyzed and classified according to their type of result – outputs, outcomes and longer-term impacts – and distributed along three different levels: individual, collective (meso), and systemic change. Insights about intercultural communication in qualitative research are discussed in relation to the characteristics of UOC researchers and the class composition of faculty and students in the Middle East and North Africa countries (MENA) region.

**Conclusions/Significance:** This analysis fills a gap in the knowledge of social impact assessment through the establishment of a new methodological approach. The course has proven to be a valuable source of social capital and provided a space to build a network of professionals with a shared focus on how to treat CL in particular – a community of practice – and increase knowledge of other skin NTDs.

**Author summary:** Neglected tropical diseases (NTDs) affecting the skin (Buruli ulcer, cutaneous leishmaniasis, post-kala-azar dermal leishmaniasis, leprosy, lymphatic filariasis, mycetoma, onchocerciasis, scabies and yaws) cause considerable disability, increase stigma and exacerbate poverty. Cutaneous leishmaniasis (CL) is one of the most neglected NTDs. It has an estimated incidence of between 900,000 and 1.3 million new cases worldwide each year. A co-venture between the World Health Organization (WHO) and Universitat Oberta de Catalunya (UOC) has developed an on-line interactive course entitled “Clinical management of CL” to provide health personnel with thorough and up-to-date information on the various aspects of CL. Using a new methodological approach, we have conducted a social impact assessment of this training program in order to identify the different type and level of results. The most significant impact has been the mobilizing of students and professors from the different editions of the course in a community of practice, thereby enhancing learning, knowledge sharing and effectivity.

## Introduction

Neglected Tropical Diseases (NTDs) are defined by the World Health Organization (WHO) as a group of 20 conditions affecting more than a billion people worldwide [1]. These conditions place an immense burden on individuals, families and communities in developing countries, thereby aggravating the cycle of poverty. Although significant improvements were achieved during the Millennium Development Goal (MDG) era, major challenges remain, including the need to reduce the impact of NTDs in endemic areas. Under the 2030 UN Agenda, the Sustainable Development Goal (SDG) 3: “Ensure healthy lives and promote well-being for all at all ages” includes the target (3.3) to end the epidemics of neglected tropical diseases by 2030 [2]. Despite this, NTDs are poorly known and rarely addressed in Medical Schools [3].

Cutaneous leishmaniasis (CL) is one of the most neglected NTDs. It has an estimated incidence of between 900,000 and 1.3 million new cases worldwide each year, and impacts the poorest populations, mainly in tropical and sub-tropical areas [4–5]. The disease is caused by the *Leishmania* parasite transmitted in an infected female sand-fly bite and the main clinical manifestations consist of cutaneous lesions that leave permanent skin scars and cause significant stigmatization [6–9].

In 2013, in a co-venture with the World Health Organization (WHO), Universitat Oberta de Catalunya (UOC) developed an on-line interactive course entitled “Clinical management of CL” to provide health personnel with thorough and up-to-date information on the various aspects of CL. Through the course, students acquired the theoretical background and “hands-on” clinical skills required for the clinical management of CL in endemic areas. The course was developed according to the WHO Manual for case management of cutaneous leishmaniasis and focused on detection and management of the disease [10].

### The Cutaneous Leishmaniasis on-line course

The e-learning course consisted of 6 ECTS (European Credit Transfer System, equivalent to 150 study hours) and was directed at clinicians, nurses, public health professionals and policy makers. Students received up-to-date information on the natural history, epidemiology, diagnosis, treatment and surveillance of the disease. The teaching strategies applied were online, sequential and participatory. Learning resources were largely based on scientific articles, WHO manuals and clinical cases and problems, as well as the study and sharing of different field experiences.

The Faculty team consisted of a UOC tutor and a Professor of Dermatology. The former was in charge of observing dynamics within the virtual classroom and specifying activities and learning strategies, guiding students, encouraging participation and facilitating learning. Student’s achievements were measured through continuous assessment activities and completed with a multiple-choice test. Once the class was completed, students were asked to provide feedback on the course through a structured questionnaire, which served to monitor the quality of the course, and improve its structure, content, and teaching methods.

With three editions of the program successfully completed we consider assessment of the social impact of the course beyond knowledge acquisition to be of paramount importance. To this end, we conducted a study to determine the impact of the CL on line course on professional practice.

### The Social Impact Assessment model

Based on several theoretical models that have been used to analyze the social impact of various interventions, mainly in the biomedical sector [11–16], the following basic considerations need to be taken into account: impacts are multifactorial, and thus the model requires a longitudinal systematic approach; the focus of the study is direct impact, therefore indirect effects are also considered but not measured; structural conditioning must be taken into account as this may present barriers or facilitators depending on context, particularly through cultural aspects; outputs (immediate effects) and deeper transformations (outcomes and impacts) must be differentiated; at the individual level, change can present in three different layers: a) knowledge acquired: cognitive and intellectual results; b) thinking level: attitudes, values, perceptions; and c) action level: behaviors and calls to action.

Social impact is considered a process rather than a situation at any given moment of time. It is a chain of results that stem from an initial input, and includes outputs (immediate results), outcomes (shortly after the intervention) and impact (global results, deeper transformations not only related to the intervention but multifactorial). Results can be observed at the individual level (micro level), the small group level (meso or interactive level) and the macro level. Multiple factors and contextual conditionings alter the course of systemic and structural change. The aim of this study is to determine the impact of the CL on-line course on professional practice

## Methods

### Population of study

Three editions of the “Clinical management of CL” course have been provided since 2014. The first two editions were taught during the 2nd semester of 2014 to two class groups, one in French and the other in English. The third edition took place during the 1st semester of 2016 in French. The course targeted healthcare professionals in MENA countries (Middle East and North Africa countries, according to grouping by the World Bank) involved in clinical care. Student enrollment was conducted following a selection process carried out by the Health Ministries of each participating country, namely Afghanistan, Algeria, Chad, Morocco, Pakistan, Syrian Arab Republic, Tunisia and Yemen, with WHO covering their expenses. The course was oriented towards health professionals, with priority given to general practitioners, nurses and dermatologists.

### Analysis methods: personal interviews

In-depth personal Interviews were used to gather information from participants. A number of topics were covered, including participants’ expectations of the course, and their actual experience during the program (Table 1 details the guidelines used for interviews). To quantify impact we considered outputs (immediate results at the end of the program) and outcomes (1 year after completing the course).

All interviews were recorded and transcribed. Our analyses were based on coded information about the different dimensions explored during the interviews, with each inquiry corresponding to at least one code. Our analysis was organized accordingly and the results section follows the same structure. Interviewee responses were compared, with their countries of origin and position (student or teaching staff) taken into consideration. The software used was Atlas.Ti v7.

**Table 1:**
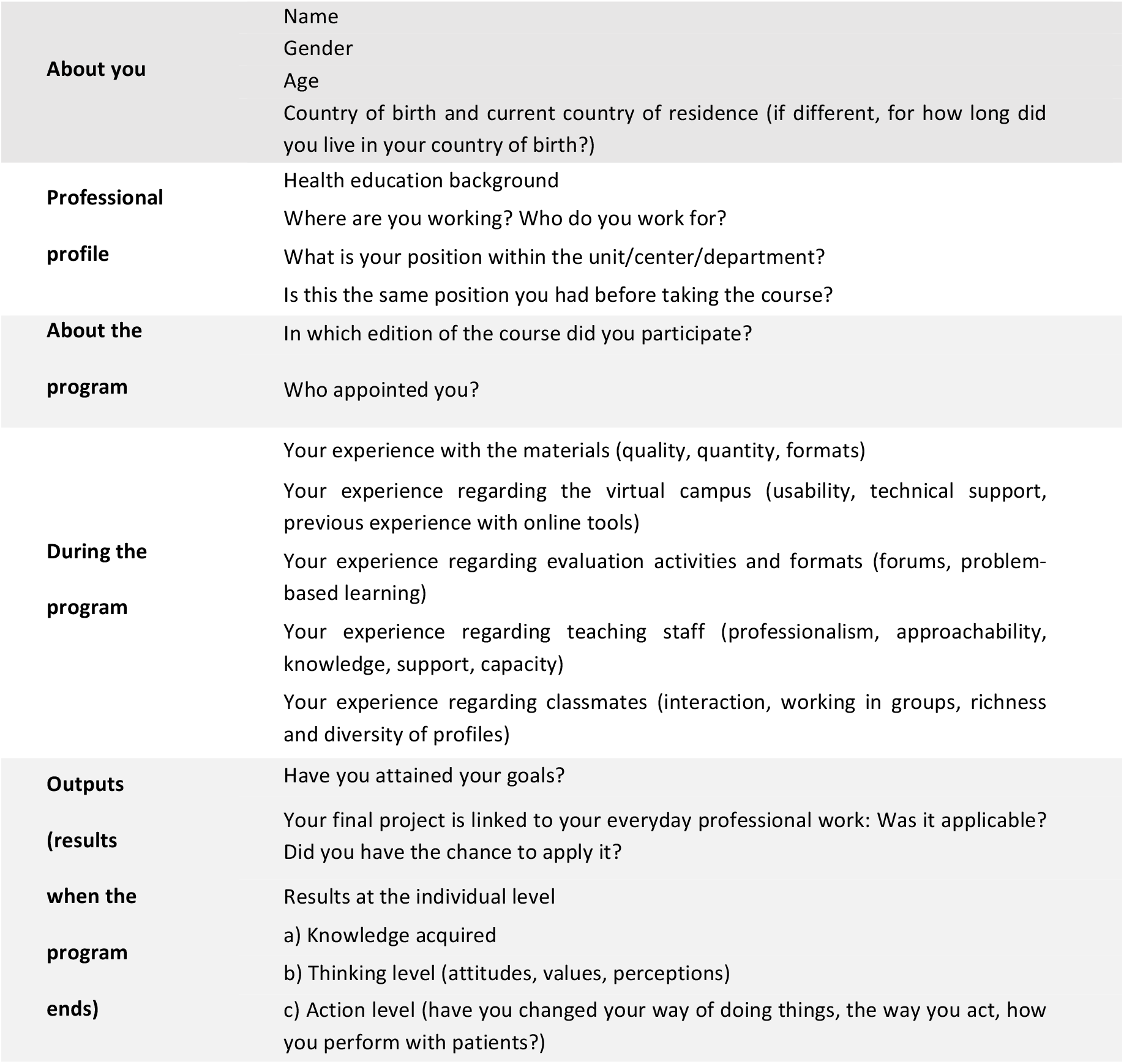
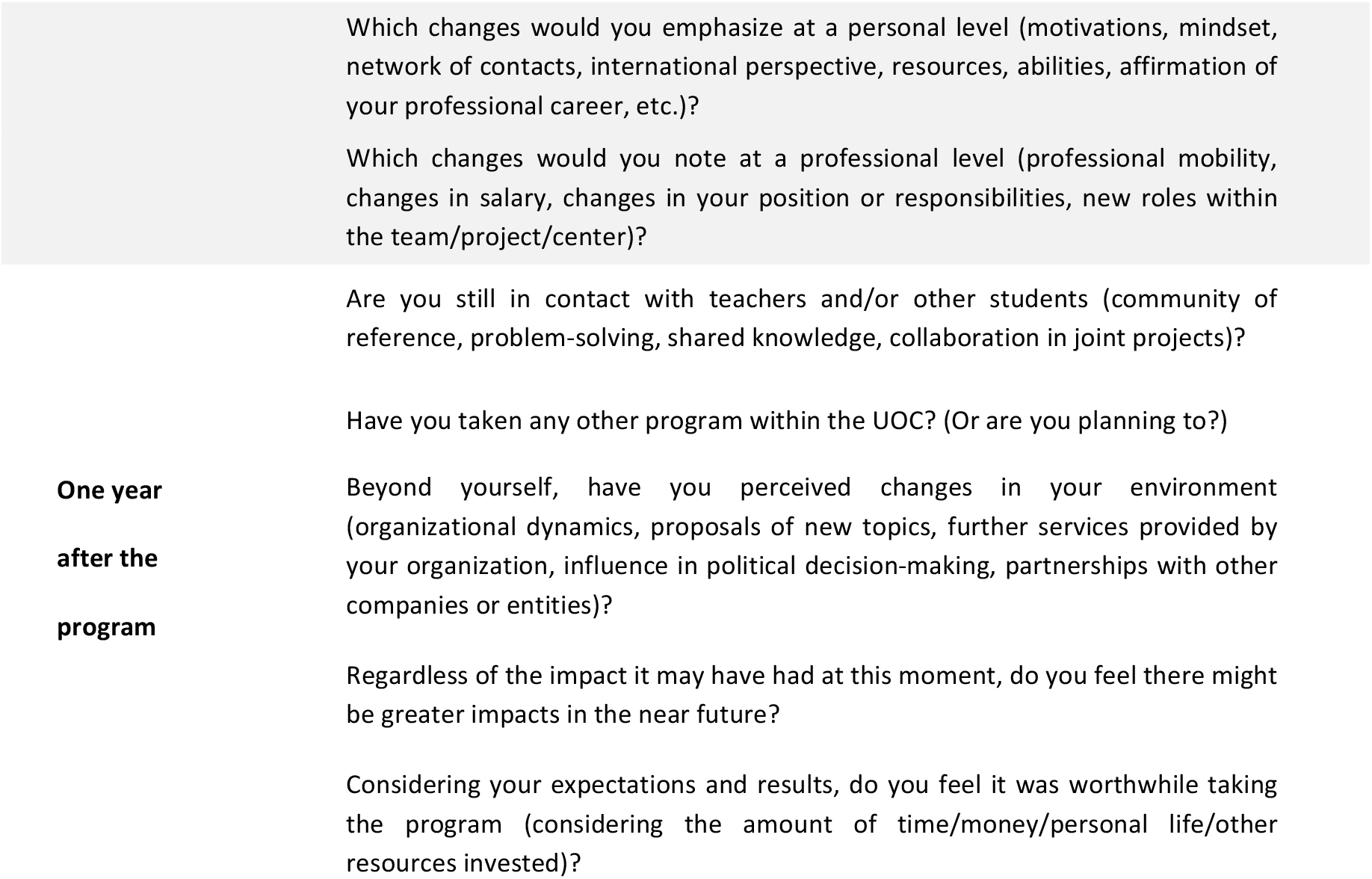
Interview guidelines (English version)

### Sample and interviewees

The eligible sample consisted of the 36 students who completed one of the three editions of the course. An institutional invitation from UOC was sent by email to these students. A number of the students who were unsure of their availability and accessibility at a certain time were invited to provide written answers to the questionnaire.

## Results

### General features of the program and the student’s universe

Sixty students from seven different countries were accepted on the three editions of the course, with the majority of participants being general practitioners. Thirty-six students (60%) succeed in passing the course. Table 2 details the number of enrollments, graduates and the drop-off rate by country. The average final score was grade B (71%) out of 3 possible grades (A, B and C).

**Table 2:** Three editions enrollment, graduates and drop-off rate by country

|  | Enrolled<br>N (%) | Graduates<br>N | Drop-off rate<br>(%) |
| --- | --- | --- | --- |
| Afghanistan | 4 (6,7) | 2 | 50,0 |
| Algeria | 9 (15,0) | 4 | 55,5 |
| Chad | 2 (3,3) | 2 | 0,0 |
| Morocco | 15 (25,0) | 11 | 26,7 |
| Syrian Arab Republic | 8 (13,3) | 4 | 50,0 |
| Tunisia | 20 (33,3) | 12 | 40,0 |
| Yemen | 2 (3,3) | 1 | 50,0 |
| **TOTAL** | 60 | 36 | 40,0 |

### Participants

Although 11 students had agreed to participate out of the eligible sample of 36, the actual number of interviewees was 7 (2 teaching staff and 5 students). Several barriers to accessing the students were identified: first, lack of feedback or interest in the request to participate, even when the program *ambassador* was the person sending the information; second, technical difficulties, including a lack of phone or internet signal, and limited skills with video chat software and virtual environments; third, logistical barriers, as a number of participants were on humanitarian missions with limited access to internet, computers and even phones.

The list of interviewees was as follows:

JP – WHO staff member and course teacher, 50-year-old male.

MM – Tunisian dermatologist and course instructor, 52-year-old male.

MI – Nurse, 35-year-old male, Morocco. Student in the 2^nd^ edition

BF – Health practitioner (16 years of experience), 45-year-old male, Tunisia. Student in the 2^nd^ edition. Recently appointed as regional Health Supervisor for the Sidi-Bouzid area.

HD – General practitioner, 39-year-old male, Morocco. Student in the 1^st^ edition. Had to abandon the program at the very beginning due to personal and family issues, but claims he would have liked to complete it.

SB – Sports Doctor, 45-year-old male, Morocco. Student in the 1^st^ edition. Head of parasitic diseases since 2014.

JB – Epidemiologist, aggregate professor in preventive health working at the preventive unit of the Pasteur Institute, 37-year-old female, Tunisia. Student in the 1^st^ edition.

### Student experience during the course

In general, students were highly satisfied with the program. They particularly emphasized the fruitfulness of the *online* discussion forums, enabling them to interact with people in different countries and environments (JP, BF, SB). The sharing of a “virtual” space online also fostered a sense of belonging and, to some extent, provided relief as they became aware of other professionals with limited knowledge of the disease.

Practical sessions based on real clinical cases proved to be the most useful, inspiring and motivating. Problem-based learning introduced significant concepts and ideas, particularly as participants had to prepare clinical cases to discuss with the others (BF). Additionally, students took part in practical sessions, which were collectively prepared with the dermatologist acting as mentor. Those students who were the most focused or most interested in scientific evidence rated this element positively, believed it was well grounded (BF), with trustworthy references and very comprehensive materials (SB).

The online format of the course was also evaluated positively. The students particularly appreciated the ability to self-administer the training content according to their own schedules. However, those who had limited computer skills suffered from the outset (BF). Moreover, it was argued that the rapid pace of the program had a negative impact on personal and family life, as a number of students had to work at night while others were sleeping: “most of those who have left the course, left due to the time-consuming design” (BF).

Participants suggested improvements to some aspects of the platform, such as simplifying it and familiarizing students through additional training with course tools prior to the course itself. A few of those students taking the French version of the program had found that some areas of the platform were in English only (JP). Although the web-based format of the class was considered to provide autonomy and be easily accessible, some students suggested the possibility of organizing “live” meetings: “At least one face-to-face session would be indispensable and promote even greater engagement with other participants” (JB).

### Outputs and outcomes

At the end of the program, all students were familiar with the CL Recommendations and Manuals, an outcome that was also emphasized and highly valued by the WHO course instructor; “knowledge acquisition regarding the disease is clear” (JP), as well as the students. The teaching staff identified vast improvements in terms of disease management awareness as the class progressed.

A key aspect was that students felt more aware, empowered and self-confident: “With this course I have learnt about diagnosis, differential diagnosis and treatment, therapeutic treatment and the subsequent follow-up; also about side-effects” (SB). According to the WHO course instructor, and as stated in WHO documents, this was extremely important in terms of recommendations on how to manage CL spreading by word of mouth, “as we make recommendations, but each country is free to adopt them or not” (JP). Providing knowledge directly to professionals appears to position those recommendations as a standard to follow, making applicability feasible and plausible. Even in terms of science planning the class proved useful; the researcher at the Pasteur Institute claimed that beyond the lessons learnt, she had broadened her research options, focus and areas of interest. In other words, according to this participant, the program may have some impact on research agendas at institutions concerned with this disease.

### Specific impacts of the program

Besides the outputs and outcomes, three levels of impact can be considered: personal, professional and systemic. While the first two remain at an individual level, the latter has wider influence.

*Personal impacts* described by the students tended to be linked to emotions. The most widespread feeling was one of increasing personal motivation for their mission and occupation (BF, SB, JB), and to a lesser extent, rising self-esteem: “To me, the best thing is the pride I feel to have been chosen. And to be working within a center which is participating in an innovative pilot program from Morocco” (MI). It is likely this pilot program made nurse MI more receptive and eager to take the course.

Regarding the *professional impact* of the course, students described their new standing as experts of reference within their teams: “Suddenly, I have become a source of information for my colleagues. Most of them come to me with questions and doubts. I have gained lots of self-confidence in diagnosis and treatments, I feel more motivated and my patients are happier. I am in the same position, but my reputation has increased a lot among my peers” (MI).

One student was appointed as a supervisor, but had probably already been selected for the post: “It was simultaneous; I was a general practitioner when I started the course, and by the end of it I was appointed Supervisor” (BF). The significance here is that he now has the ability to disseminate the knowledge he has acquired: “In fact, now I supervise my colleagues I can share my knowledge on the disease with them and their patients” (BF).

Finally, and importantly, the course has proven to be a valuable source of social capital. Students are highly satisfied with the course and still in contact with the teaching staff (mainly the Tunisian dermatologist) and each other for access to further information about topics relating to several skin NTDs (MM). In other words, the program has provided a space to build a network of professionals with a shared focus on how to treat this skin NTD in particular, and increase knowledge of other skin NTDs. This sense of a community of practice and being part of a privileged elite may also strengthen application of the concepts and methods learnt during the course. At the institutional level, a highly positive outcome was an expanded program on skin NTDs provided in 2017 and 2018, again by UOC and WHO, covering Buruli ulcer, cutaneous leishmaniasis, leprosy and yaws.

## Discussion

The course has proven to be a good source of social capital. As stated above, a network of professionals with a shared focus has been created, and students still contact the teaching staff and each other for further information about topics relating to several skin NTDs. This sense of a community of practice and belonging to a privileged elite may also reinforce application of the concepts and methods learnt during the course. At the end of the program all the students had become familiar with WHO recommendations. A key aspect is that students are not only more aware, they feel empowered and self-confident; this is extremely important for the dissemination of guidelines. Providing knowledge directly to professionals appears to position the recommendations as a standard to be followed, making applicability feasible and plausible.

Personal impacts described by the students tended to be linked to emotions. The most widespread feeling was one of increasing personal motivation for their mission and occupation, and to a lesser extent, rising self-esteem. Regarding their professional performance, the students have gained a new standing as experts of reference within their teams.

To improve and increase the potentialities of interventions it is crucial to analyze their social impact. Although this process of analysis has provided limited reliable knowledge, a number of discoveries have been made about the challenges of evaluating a program of this nature. Our initiative to assess the impact of the course was an instrumental program: it was short-lasting and oriented to a very specific type of knowledge acquisition. In terms of impact, it would have been constructive to gain access to statistics regarding the number of patients diagnosed and treated according to the knowledge acquired on the course, although this would be influenced by the national guidelines of each country, which are not always aligned with WHO recommendations.

The scope of this assessment is somewhat limited due to the methodology chosen, with only seven of the 36 students able to take part in the interviews. Alternative methods, such on-site focus groups, would have more effectively served our purpose, but this was not possible. It is also important to keep in mind that the course is targeted at health professionals established and practicing in MENA countries, while the impact assessment was carried out by European researchers. To minimize bias, three important aspects should be addressed: the digital divide affecting some of the participants, cultural differences, and the importance of providing feedback to a stranger (the interviewer).

### Limitations

Potential impacts are difficult to measure in part due to intrinsic work dynamics and the high turn-over and instability of healthcare employees from institutions in the targeted countries. These are structural factors that may slow down dissemination and limit the possibilities of bringing the expertise acquired into the field. Once the course was completed there was no follow-up or space for continuous interaction with the faculty. Those involved would prefer to maintain some communication flow as the course has not only been a space for learning, but also one in which trust in the instructors has been established.

### Conclusions

The course has proven to be a valuable source of social capital. Students are highly satisfied with the course and still contact the teaching staff and each other for further information about topics relating to several skin NTDs. In other words, the program has provided a space to build a network of professionals with a shared focus on how to treat this skin NTD in particular, and increase knowledge of other skin NTDs. This sense of a community of practice and being part of a privileged elite may also strengthen application of the concepts and methods learnt during the course.

## References

1. World Health Organization. Neglected Tropical Diseases. Available from: http://www.who.int/neglected_diseases/diseases/en/. Accessed April 17, 2019.

2. United Nations. Sustainable Development Goals. Available from: https://www.un.org/sustainabledevelopment/sustainable-development-goals/ Accessed April 17, 2019.

3. World Health Organization. Integrating neglected tropical diseases into global health and development: fourth WHO report on neglected tropical diseases. 2017. ISBN 978-92-4-156544-8.

4. Alvar J, Vélez ID, Bern C, Herrero M, Desjeux P, Cano J, Jannin J, den Boer M, WHO Leishmaniasis Control Team. Leishmaniasis worldwide and global estimates of its incidence. PLoS One. 2012;7(5): e35671. Doi: 10.1371/journal.pone.0035671.

5. Bailey F, Mondragon-Shem K, Hotez P, Ruiz-Postigo JA, Al-Salem W, Acosta-Serrano Á, Molyneux DH. A new perspective on cutaneous leishmaniasis-Implications for global prevalence and burden of disease estimates. PLoS Negl Trop Dis. 2017;11(8): e0005739. Doi: 10.1371/journal.pntd.0005739.

6. Khatami A, Emmelin M, Talaee R, Miramin-Mohammadi A, Aghazadeh N, Firooz A, Stenberg B. Lived Experiences of Patients Suffering from Acute Old World Cutaneous Leishmaniasis: A Qualitative Content Analysis Study from Iran. J Arthropod Borne Dis. 2018;12(2): 180–195.

7. Bennis I, De Brouwere V, Belrhiti Z, Sahibi H, Boelaert M. Psychosocial burden of localised cutaneous Leishmaniasis: a scoping review. BMC Public Health. 2018;18(1): 358. Doi: 10.1186/s12889-018-5260-9.

8. Bennis I, Belaid L, De Brouwere V, Filali H, Sahibi H, Boelaert M. “The mosquitoes that destroy your face”. Social impact of Cutaneous Leishmaniasis in South-eastern Morocco, a qualitative study. PLoS One. 2017;12(12): e0189906. Doi: 10.1371/journal.pone.0189906. eCollection 2017.

9. Bailey F, Mondragon-Shem K, Haines LR, Olabi A, Alorfi A, Ruiz-Postigo JA, Alvar J, Hotez P, ER Adams, Vélez ID, Al-Salem W, Eaton J, Acosta-Serrano A, Molyneux DH. Cutaneous leishmaniasis and co-morbid major depressive disorder: A systematic review with burden estimates. PLoS Negl Trop Dis. 2019;13(2): e0007092. Doi: 10.1371/journal.pntd.0007092. eCollection 2019.

10. World Health Organization, Regional Office for the Eastern Mediterranean. Manual for case management of cutaneous leishmaniasis in the WHO Eastern Mediterranean Region. 2014. http://www.who.int/iris/handle/10665/120002

11. Adam P, Solans-Domènech M, Pons JMV, Aymerich M, Berra S, Guillamon I, Sánchez E, Permanyer-Miralda G. Assessment of the impact of a clinical and health services research call in Catalonia. Res Eval. 2012;21(4): 319–328.

12. Brook S, Thomson A, Strode M. DFID New Economists Guide: Monitoring Chapter. Oxford: Oxford Policy Management; 2006.

13. Buxton M, Hanney S. How can payback from health services research be assessed? Health Serv Res Policy. 1996;1(1): 35–43.

14. De Vries H, Kremers S, Smeets T, Brug J, Eijmael K. The effectiveness of tailored feedback and action plans in an intervention addressing multiple health behaviors. Am J Health Promot. 2008;22(6): 417–425.

15. Forsetlund L, Bjørndal A, Rashidian A, Jamtvedt G, O’Brien MA, Wolf FM, Davis D, Odgaard-Jensen J, Oxman AD. Continuing education meetings and workshops: Effects on professional practice and health care outcomes. Cochrane Database of Systematic Reviews 2009. Doi: 10.1002/14651858.CD003030.pub2

16. Spencer R. Nurses’, midwives’ and health visitors’ perceptions of the impact of higher education on practice. Nurse Educ Today. 2006;26(1): 45–53.

